# How Prior Motor States Can Shape Perceptual Decision Bias: Insights from Sensorimotor Beta Oscillations

**DOI:** 10.1101/2025.07.21.665769

**Authors:** Ningjing Cui, Robert J. van Beers, Jeroen B. J. Smeets, Bernadette C. M. van Wijk

## Abstract

Emerging evidence suggests that decision making is not a purely cognitive process preceding motor output but rather an embodied phenomenon in which the sensorimotor system actively shapes choice selection. Beta oscillations (13–30 Hz) over left and right sensorimotor cortex have been shown to encode motor action and exert inhibitory influence through lateralization of the post-movement rebound, biasing subsequent decisions toward responses with the other hand. This study investigated whether sensorimotor beta oscillations elicited by an isolated choice-unrelated motor action could influence a subsequent perceptual decision. Twenty-nine healthy adults (25 right-handed) completed a two-stage task while 64-channel EEG was recorded. In each trial, participants executed an initial button press with either the left or right thumb to a letter cue (“L” or “R”) followed by, after a variable delay (0.5, 1.2 or 3 s), a briefly presented visual grating requiring a decision response on the orientation of the grating via a second button press. We computed the degree of beta power lateralization between left and right (pre-) motor cortical sources that was present during presentation of the grating stimulus and quantified choice bias as the threshold difference between psychometric curves fitted to left-versus right-initial response trials. Participants with stronger beta lateralization during stimulus presentation showed a greater tendency to alternate their decision response from the hand used for the initial button press and exhibited slower decision speeds. However, we failed to detect a significant group-level decision bias induced by the initial motor action, nor were there consistent within-subject correlations between decision bias and beta lateralization across delays. Following the perspective of embodied-decision making, our results partially support the influence of motor cortex activity on the choice process, while also suggesting that beta activity may serve as a trait-like index of individual susceptibility to decision bias.

## 1. Introduction

Perceptual decision making describes the process by which sensory information from the environment is transformed into goal-directed actions in both humans and animals (Cisek & Pastor-Bernier, 2014; Gold & Shadlen, 2007; Heekeren et al., 2008). It is commonly investigated using paradigms such as motion discrimination (e.g., Hanks et al., 2006; Huk & Shadlen, 2005) and orientation judgment tasks (e.g., Assarioti et al., 2025; Zhang et al., 2019), where participants make binary choices by selecting motor responses that reflect stimulus-related decisions. Traditionally, perceptual decision making has been characterized as a serial process (Lepora & Pezzulo, 2015). This process starts with sensory evidence being encoded in the corresponding sensory cortex (Britten et al., 1996; Newsome & Pare, 1988). The evidence is then integrated in association regions such as posterior parietal and prefrontal cortex (Huk & Shadlen, 2005; Zhou & Freedman, 2019). Finally, the resulting decision signals are transmitted to sensorimotor areas such as the premotor cortex for action execution (Cisek & Kalaska, 2005). This framework suggests a functional dissociation between cognitive processing circuits (i.e., posterior parietal and prefrontal cortex) where decisions are thought to be formed, and the sensorimotor system which is considered solely as the final output stage. However, accumulating neurophysiological evidence challenges this view by showing that competing choice-related action plans are concurrently activated within sensorimotor areas even prior to the final commitment of a choice in higher-order cognitive regions (Cisek & Kalaska, 2005, 2010; Pastor-Bernier & Cisek, 2011). These findings support a parallel processing model, where choice formation and action specification unfold simultaneously and are dynamically intertwined (Lepora & Pezzulo, 2015).

In contrast to traditional lab-based experiments, decision making in real-word scenarios often occurs in dynamically changing environments where costs and affordances of possible actions actively shape the process of choice formation (Cisek, 2007). This idea has led to the embodied decision-making framework (Cisek & Pastor-Bernier, 2014; Lepora & Pezzulo, 2015), which posits that the sensorimotor system is not merely a passive implementer of decisions but is actively involved in and even capable of influencing the decision making itself. For instance, in a predator-prey interaction scenario, a lion choosing between two moving preys may continuously adjust its preference based on fluctuating factors such as movement trajectory and distance, demonstrating how motor affordances shape choice commitments over time. A growing body of behavioural evidence has indicated that motor-related factors influence the choice formation. For example, using computational modelling, Lepora and Pezzulo (2015) found that incorporating a “commitment” parameter that captures a bias toward an initially selected option, even as accumulating evidence starts to favour the alternative, provides a better fit to empirical data. This indicates that the dynamics of an unfolding action can bias the ultimate choice. Additional evidence comes from observations demonstrating that an increased motor effort of the response, such as elevated physical resistance (Burk et al., 2014; Hagura et al., 2017) or greater greater reaching distance (Assarioti et al., 2025; Marcos et al., 2015) biases choice behaviour toward the less effortful option.

The embodied decision-making framework also offers a different perspective on the phenomenon that current choices are not only determined by ongoing sensory input, but also biased by previous choices. Inherently, previous choices are accompanied by a motor response to indicate the selected choice, which leaves the possibility that it might be the motor response per se, rather than the choice itself, that influences subsequent decisions. Behavioural studies examining how previous motor responses affect choice bias have yielded mixed findings. Some studies found no significant contribution of motor history (Fernberger, 1920; Akaishi et al., 2014; Braun et al., 2018) while others reported clear motor-history effects (Pape et al., 2017; Urai et al., 2017; Zhang and Alais, 2020). These inconsistencies may be attributed to differences in task design, such as the presence or absence of feedback (Fründ et al., 2014), or the use of variable delays between motor response and decision cues. Most studies adopted a varied choice-response mapping paradigm, as in the study by Zhang & Alais (2020) where identical perceptual choices were randomly assigned to different motor responses, explicitly dissociating the contribution of motor history from that of perceptual choice. They found an alternating motor bias, where participants were inclined to alternate motor responses, regardless of the associated perceptual choice. Similarly, Pape et al. (2017) showed that a choice-unrelated motor action preceding a decision task reduced choice repetition, further suggesting the direct influence of prior motor responses on subsequent decisions.

Neurophysiologically, a motor history bias might arise through the temporal dynamics of sensorimotor beta oscillations that occur around each movement. Beta oscillations refer to rhythmic activity in the 12 to 30 Hz frequency range and are widely considered as a neural index of motor preparation and execution (Jurkiewicz et al., 2006; Kilavik et al., 2013; Pfurtscheller & Da Silva, 1999; van Wijk et al., 2012). During brief movements, a reduction in beta power (i.e., beta desynchronization) is typically observed starting around 0.5 seconds before movement onset and continuing throughout movement execution (Leocani et al., 1997; Takemi et al., 2013). This is followed by a rapid increase in beta power (i.e., beta rebound), beginning approximately 0.5 seconds after movement offset and lasting up to 4-5 s in some cases (Bailey & Bardouille, 2025; Houdayer et al., 2006). Beta desynchronization is thought to reflect active motor preparation, whereas the subsequent beta rebound has been linked to processes that inhibit motor output (Jurkiewicz et al., 2006; Pfurtscheller et al., 1996; Van Wijk et al., 2009). This is supported by findings of slower responses when actions are initiated during the rebound phase (Muralidharan & Aron, 2021; Zhang et al., 2024) and reduced corticospinal excitability during this period (Chen et al., 1998). Crucially, the beta rebound is typically stronger in the sensorimotor areas contralateral to the moving hand compared to the ipsilateral side (Jurkiewicz et al., 2006; Pfurtscheller et al., 1996). This lateralization can hypothetically lead to a bias in subsequent decisions, especially when new stimulus information is presented while the beta rebound triggered by the previous motor response is still ongoing.

Two MEG studies have directly examined motor history effects in perceptual decision making through the lens of sensorimotor beta oscillations (Pape & Siegel, 2016; Urai & Donner, 2022). Pape and Siegel (2016) used a random dot motion task with a varied choice-response mapping paradigm that allowed them to dissociate the influence of previous responses from choices. Their results indicated that a stronger lateralized beta rebound was associated with a higher likelihood of choice alternation, both at the single trial level and across participants. By contrast, Urai and Donner’s (2022) study did not indicate a difference in sensorimotor beta lateralization during stimulus presentation between participants who had a tendency to repeat their previous motor response and those who tended to alternate. Yet, using drift-diffusion modelling, the authors found that the degree of beta power lateralization internally biased the starting point of evidence accumulation toward alternation, hence still indicating an influence on the decision-making process. Both studies employed relatively complex behavioral paradigms, involving choice-response mappings, delayed response requirements, and, in the case of Urai and Donner (2022), trial-by-trial performance feedback. These features may have introduced additional cognitive demands or strategies, which could have potentially confounded the effects of motor history. Some studies (Donner et al., 2009; de Lange et al., 2013), although not explicitly testing motor history effects, employed a fixed stimulus-response mapping and found that spontaneous fluctuations in beta lateralization could partially predict subsequent choices, suggesting a potential top-down regulatory role of the motor cortex in decision-making.

To investigate how prior motor states induced by an isolated, choice-unrelated motor action can influence a subsequent perceptual choice, we adopted a simple paradigm similar to Pape et al. (2017) in which a perceptual decision-making task was embedded immediately following an initial choice-unrelated movement. To ensure that this movement did not trigger additional cognitive processing, its direction was held constant within each block rather than being determined by a choice-response mapping. To probe how the evolving lateralized beta rebound shapes choice bias, we introduced three stimulus onset delays after movement termination: short (0.5 s), medium (1.2 s), and long (3 s). These time points were selected to roughly capture distinct phases of the beta rebound, with the short delay corresponding to its onset, the medium delay approximating the expected peak, and the long delay approaching a return to baseline, based on previous studies (Jurkiewicz et al., 2006; Pfurtscheller & Lopes da Silva, 1999). Considering that rebound timing varies across individuals (Cheyne et al., 2003; Jurkiewicz et al., 2006), we treated these delays as exploratory sampling points for beta dynamics. We hypothesized that sensorimotor beta lateralization around decision onset reflects an inhibitory influence on actions performed with the hand that just executed the choice-unrelated movement and would therefore drive subsequent decisions towards alternation. Accordingly, we predicted that participants with stronger average beta lateralization would exhibit larger alternation bias overall. Additionally, we expected that within each participant, delays that evoke stronger beta lateralization would exhibit higher alternation bias.

## 2. Methods

### 2.1 Participants

Twenty-nine healthy adults were recruited via the university’s online subject pool for participation in the study. All participants had normal or corrected-to-normal vision, reported no history or current diagnosis of neurological, psychiatric, or musculoskeletal disorders, and did not use sleep-inducing medications, sedatives such as benzodiazepines, or anticonvulsants such as carbamazepine. Six participants were excluded from further analyses based on visual inspection of their individual psychometric curves (see Section 2.5.1). The final sample comprised 23 participants (17 females; median age = 20.5 years, age range = 18–28 years), of whom 21 were right-handed and 2 were left-handed. All participants provided written informed consent prior to participation and were compensated with course credits. The study protocol was approved by the Scientific and Ethical Review Board of the Faculty of Behavioural and Movement Sciences at Vrije Universiteit Amsterdam.

### 2.2 Stimuli

The experiment was programmed in MATLAB R2023b (The MathWorks, Inc.). Stimuli were generated and presented using the Psychophysics Toolbox Version 3.0.14 (Brainard, 1997; Pelli, 1997; Kleiner et al., 2007). All stimuli were presented on a mid-grey background (RGB: [127, 127, 127]) using an ASUS ROG Strix XG248Q monitor (23.8″ diagonal, 1920 × 1080 resolution, 240 Hz refresh rate; physical dimensions: 526 mm × 296 mm), centrally positioned for stimulus display. A fixation cross (25 pixels in size, 8 pixels in thickness), letter cues (“L” or “R”; 40-pixel font size, Times font), and task instructions presented at the beginning of each block (40-pixel font size, Times font) were all displayed centrally in white (RGB: [255, 255, 255]).

Orientation grating stimuli (**Figure 1**) were generated using a 400 × 400 pixel sinusoidal grating. The grating had a Michelson contrast of 0.8, a spatial frequency of 0.1 cycles per pixel, and a phase of 0°, and was rendered with alpha blending and the smoothstep smoothing method to ensure gradual transitions at the aperture boundary. To increase perceptual difficulty, the grating was overlaid with a Gaussian noise texture, generated by sampling pixel intensity values from a zero-mean Gaussian distribution and applying a spatial Gaussian filter with a standard deviation of 80 pixels. The noise texture was spatially constrained by a circular alpha mask (radius = 200 pixels) to match the size of the grating aperture. A central grey annulus (radius = 40 pixels) was superimposed on the grating, allowing the fixation cross to remain visible at the centre to maintain spatial consistency throughout each trial. The orientation of the grating differed between trials.

**Figure 1.**
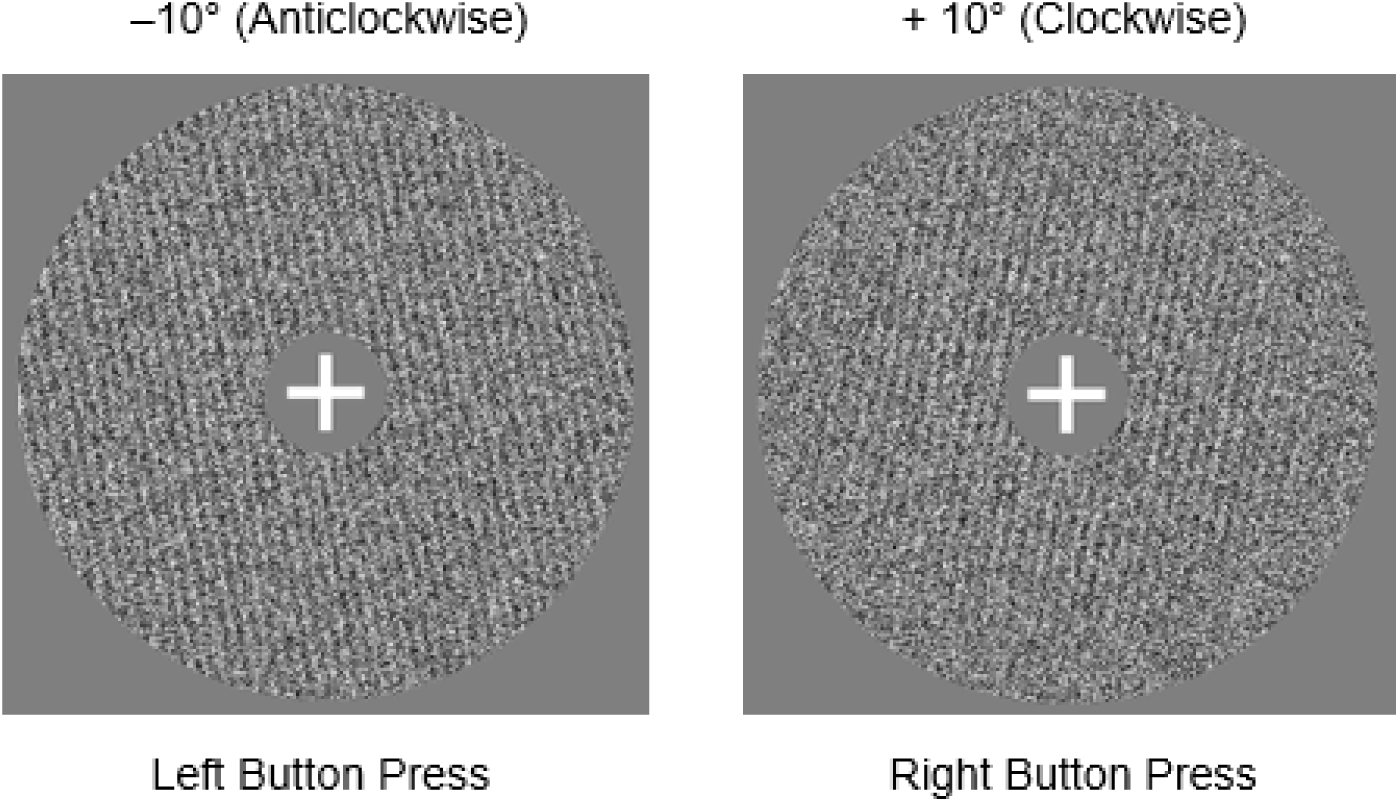
Two example orientation grating stimuli. The stimuli shown correspond to the initial orientations used in the staircase: –10° (anticlockwise, left panel), and +10° (clockwise, right panel). Below each stimulus, the correct button press is indicated.

### 2.3 Experimental Procedure and Design

The experiment was conducted in a quiet room with dim, constant artificial lighting. The experimenter was present in a separate room next door. Participants were seated in an adjustable chair behind a table with a computer monitor at a viewing distance of approximately 80 cm, wearing an EEG cap (**Figure 2**). They were instructed to hold a response box with their left and right thumbs resting on the corresponding buttons while their forearms were positioned comfortably on the table, and to respond by pressing the appropriate button according to the task requirements. The chair height was adjustable to ensure physical comfort and optimal visibility of the display. The full experimental session lasted approximately 90 minutes, comprising roughly 45 minutes of preparation time (including instructions and EEG setup) and 45 minutes of task execution, with short breaks provided between blocks to minimise fatigue.

**Figure 2.**
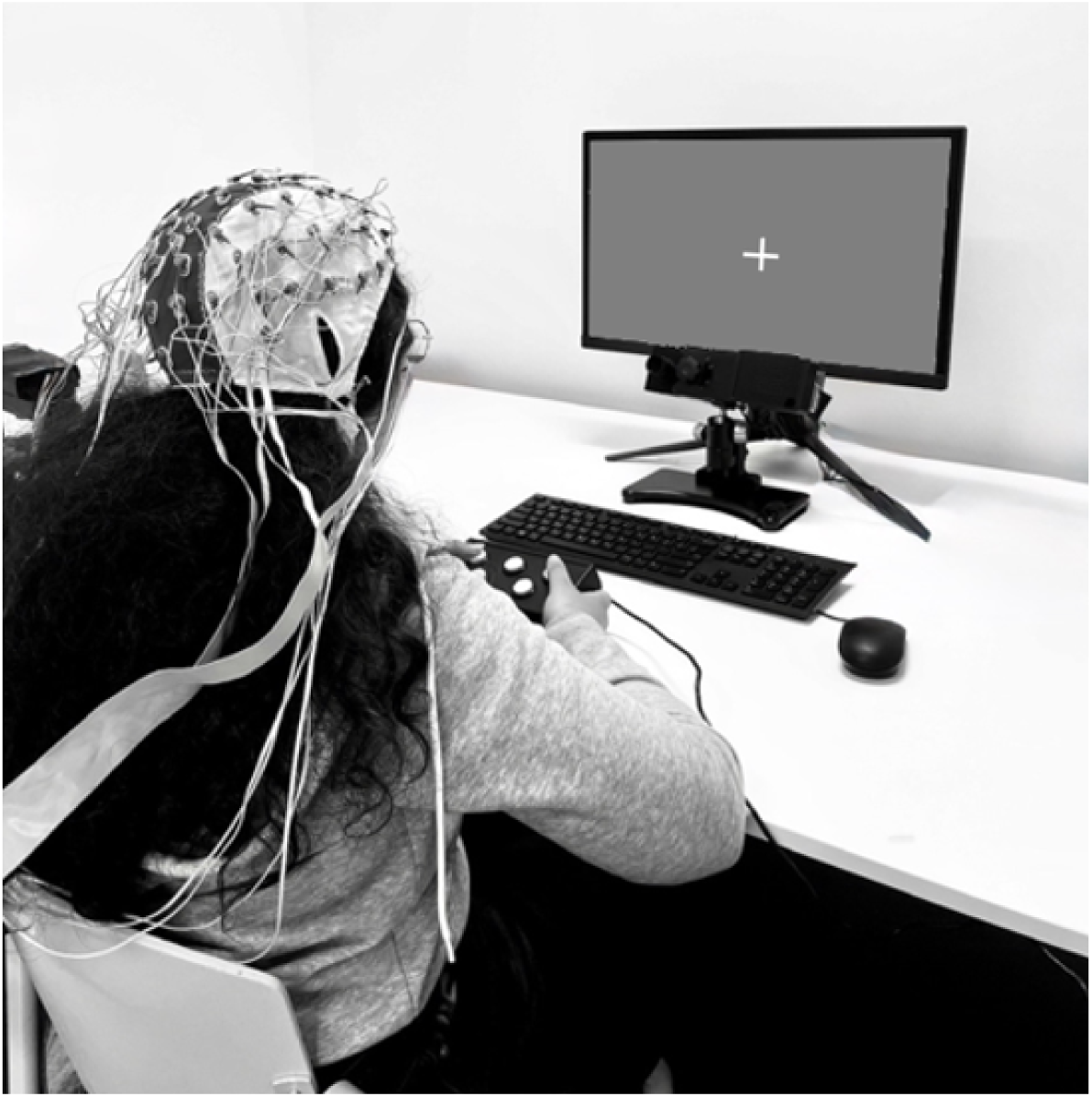
Experimental Setup. Participants were seated comfortably in front of a monitor while wearing an EEG cap. They held a response box with their left and right thumbs positioned on the corresponding buttons.

Each experimental trial consisted of two sequential stimuli: a letter cue “L” or “R” that prompted the participant to press the corresponding button with their left or right thumb, followed by an orientation grating stimulus pointing either anti-clockwise or clockwise. We will refer to the first button press as “initial response”, which merely served to induce a post-movement beta rebound in sensorimotor cortices that would still persist at the time of the grating stimulus presentation. The cue for the initial response was always the same within a block of 120 trials, so it did not require a decision. To reinforce this structure, participants received an on-screen instruction before each block indicating the required initial response (e.g., “*Within this block, you will see a letter “L” at the beginning of each trial, so please press the LEFT button to start the trial.*”). The second button press, hereafter referred to as the “decision response”, indicated the participant’s judgment of the orientation of the grating stimulus. Participants were instructed to press the left button if they perceived the grating to be tilted anticlockwise, and the right button if it was tilted clockwise (see **Figure 1** for example stimuli).

As illustrated in **Figure 3**, each trial began with the presentation of a central fixation cross displayed for 2 s, followed by a letter cue (“L” or “R”) shown at the centre of the screen. Participants were instructed to press the corresponding button (left or right) as quickly as possible. Upon release of the button, a fixation cross reappeared at the centre of the screen for one of three possible stimulus onset delays (short = 0.5 s, medium = 1.2 s, or long = 3 s), which served to ensure between-trial variability in the degree of beta rebound lateralization. Following this delay, an oriented grating appeared around the fixation cross for 0.25 s. Participants were instructed to make their decision response as quickly and accurately as possible. After the grating offset, the fixation cross remained visible for up to 2.75 s, yielding a maximum response window of 3 s. The fixation cross disappeared immediately after a response was registered, followed by a 1-s blank screen. If participants responded before the grating disappeared, the response was recorded, but the grating remained on-screen for the full 0.25 s. In this case, the trial transitioned directly to a 1-s blank screen. If no response was made within the 3-s window, the trial was recorded as a non-decision response, followed by the same 1-s blank screen. Participants completed a total of 480 trials, divided into four blocks of 120 trials each, with block order fully randomized across participants. Two blocks required left-hand initiation (i.e., the letter cue “L” presented on every trial), and two blocks required right-hand initiation (letter cue “R” presented). This resulted in a 2 (initial response: left/right) × 3 (stimulus onset delay: short/medium/long) factorial design.

**Figure 3.**
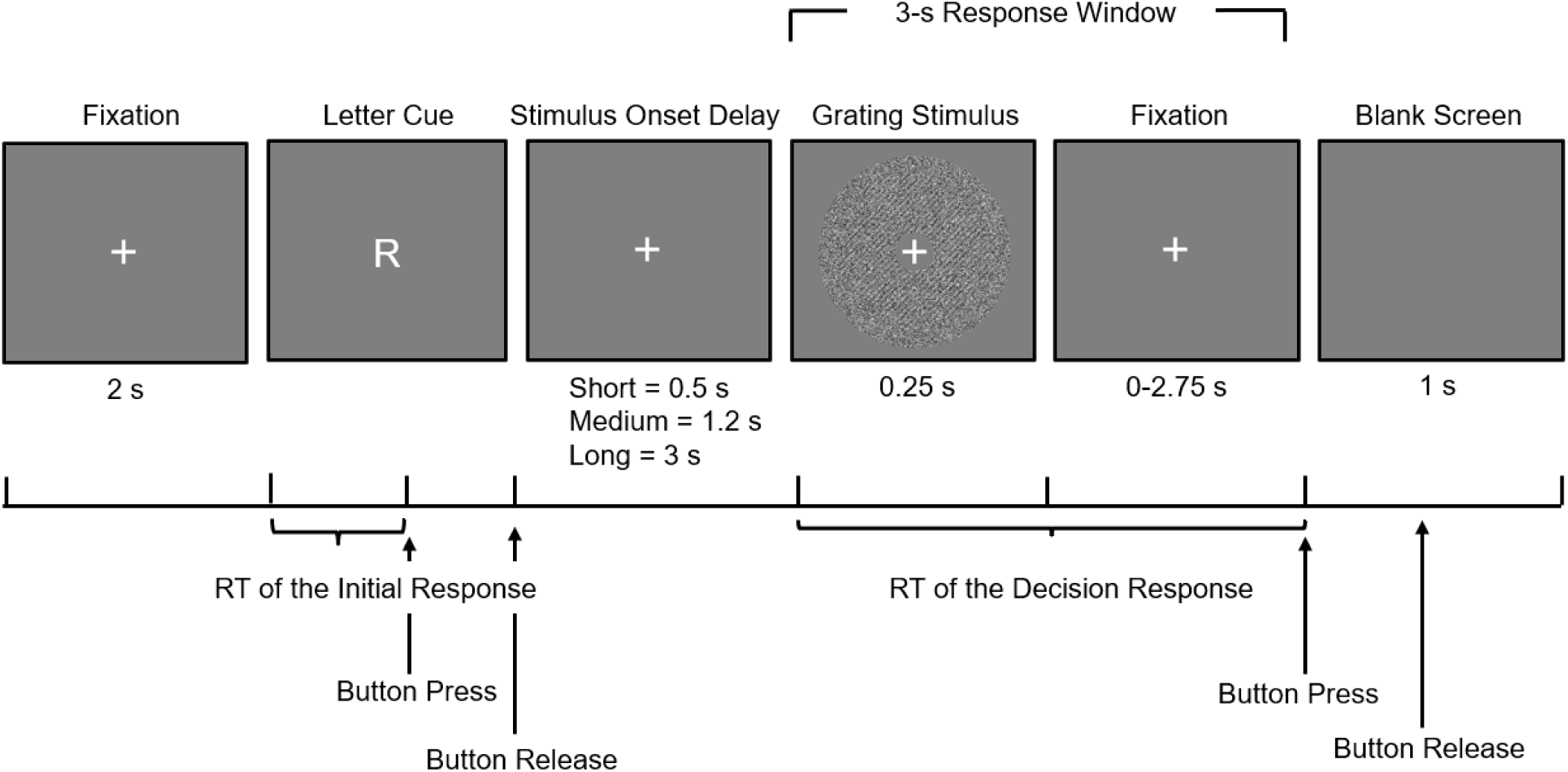
Trial sequence of the experiment. Each trial began with the presentation of a central white fixation cross displayed for 2 s, followed by a letter cue (“L” or “R”) prompting a left or right button press (i.e., initial response) (*In the illustrated example, the required response is a right-button press.*) RT of the initial response refers to the reaction time between the onset of the letter cue and the participant’s button press. Upon release of the button, the fixation cross reappeared for one of three possible delays (0.5, 1.2, or 3 s), followed by a 0.25-s decision grating stimulus. Participants responded based on perceived orientation (anticlockwise = left, clockwise = right). RT of the decision response refers to the reaction time between the onset of the decision grating stimulus and the decision-related button press. The total response window was 3 s. See main text for further details.

To maintain task difficulty, a staircase procedure (Cornsweet, 1962) was implemented within each block to determine the orientation of the grating. Each of the three stimulus onset delays (short, medium, long) had two staircases, one in which the grating was initially tilted +10° (clockwise) and one in which it was initially tilted –10° (anticlockwise) relative to vertical, resulting in a total of six independent staircase sequences per block (2 initial orientation × 3 stimulus onset delays). The six staircases were randomly assigned and equally distributed (40 trials per delay) within each block. Each staircase followed a 1-up-1-down rule based on participants’ decision responses. If a participant made a left decision response on a given trial, the orientation for the next trial in that staircase increased by 1°, shifting further in the right (clockwise) direction. In contrast, a right decision response resulted in a 1° decrease in the angle, shifting the orientation towards the left (anticlockwise) direction. For example, following a right response to a +10° (clockwise) stimulus, the next stimulus in that condition would be presented at +9°. If the response had instead been left, the next stimulus would be presented at +11°. All orientations were theoretically constrained within a range of –45° to +45° to prevent excessive deviation from vertical. In practice, orientations used in the task varied only between –12° to +12°.

Before the main experiment began, participants completed a practice block to familiarise themselves with the task. The practice block consisted of 20 trials, with the initial letter cue (“L” or “R”) constant, randomly assigned at the beginning of the block. Grating orientations followed a fixed sequence: ±45°, ±40°, ±35°, ±30°, ±25°, ±20°, ±15°, ±10°, ±5°, ±1°, with positive or negative signs randomly assigned on each trial. For the practice block only, participants received immediate visual feedback at the centre of the screen following each decision response: a green “*Correct!*” for accurate responses, a red “*Incorrect!*” for errors, and a white “*Too slow!*” if no response was made within the maximum 3-s response window. Participants were instructed to respond as quickly as possible, to attend to the variable delay between the initial response and stimulus onset, and to withhold their decision response until the grating appeared. Participants were also instructed to sit still, focus on the fixation cross, and blink only during the blank screen periods.

### 2.4 EEG Acquisition

Electroencephalography (EEG) data were recorded using a 64-channel BioSemi ActiveTwo system at a sampling rate of 2048 Hz. Electrodes were positioned according to the international 10–10 system, covering the entire scalp. During EEG acquisition, signals were referenced to the common mode sense (CMS) and driven right leg (DRL) electrodes, with scalp electrode-skin impedances kept below 10 kΩ. Offline, EEG data were re-referenced to the average signal across all electrodes. Additionally, eight external electromyographic electrodes were used to record hand muscle activity for potential validation of motor onset timing but these signals were not analysed for the current study. Event synchronization was achieved via digital triggers sent from the stimulus computer to the EEG recording system to mark trial events.

### 2.5 Data Analysis

#### 2.5.1 Behavioural Analysis

All behaviour analyses were conducted using MATLAB R2023b (The MathWorks, Inc.) and R (version 4.4.1; R Core Team). Parametric or non-parametric tests were applied depending on the normality of the data. Reaction time (RT) of the initial response refers to the reaction time between the onset of the letter cue and the participant’s button press; RT of the decision response refers to the reaction time between the onset of the decision grating stimulus and the decision-related button press (see also in Figure 2). Prior to all data analyses, we excluded trials based on the following criteria: (1) an incorrect initial response (i.e., a right-button press following the cue “L” or a left-button press following “R”); (2) an RT of the initial response exceeding 1.5 s; or (3) no registered decision response within the 3-s response window; to maintain consistency with the EEG preprocessing pipeline (see Section 2.4.2.1) and to ensure proper task engagement. A total of 159 trials (1.4% of all trials) were excluded based on these criteria, with a mean of 8.3 trials per participant (SD = 7.1).

Next, we fitted psychometric curves using *psignifit* version 4.0, a MATLAB toolbox for psychometric function estimation that fits a parametric, unidimensional sigmoid to binary responses to estimate the probability of a particular response as a function of stimulus intensity (Fründ et al., 2011; Wichmann & Hill, 2001). Psychometric fitting was initially performed on each participant’s decision responses to visually assess decision-making performance and to identify participants with insufficient task engagement, who were subsequently excluded from further analyses. Psychometric curves were fitted separately for each combination of initial response (left/right) and delay (short/medium/long) within each participant, modelling the proportion of rightward choices as a function of the grating’s orientation. Each psychometric function was modelled as a cumulative Gaussian, with all decision responses equally weighted. From each fit, we extracted three parameters: (1) the threshold, representing the orientation at which 50% of responses were rightward; (2) the width, defined as the distance between the 5th and 95th percentiles of the unscaled cumulative Gaussian, reflecting the participant’s sensitivity to changes in orientation; (3) the lapse rate, representing the proportion of stimulus-independent responses and corresponding to the upper and lower asymptotes of the fitted curve. We excluded participants with poor task engagement and unreliable perceptual performance. Specifically, participants were excluded if their mean lapse rate across all six combinations exceeded 7%, or if any individual psychometric curve exhibited a lapse rate greater than 15%. Additionally, we carefully inspected each psychometric fit to ensure that the data points did not show any considerable deviations from the fitted curve. Based on these criteria, six participants were excluded, resulting in a final sample of 23 participants included in subsequent analyses (See **Figure 4** for examples of good and poor task performance).

**Figure 4.**
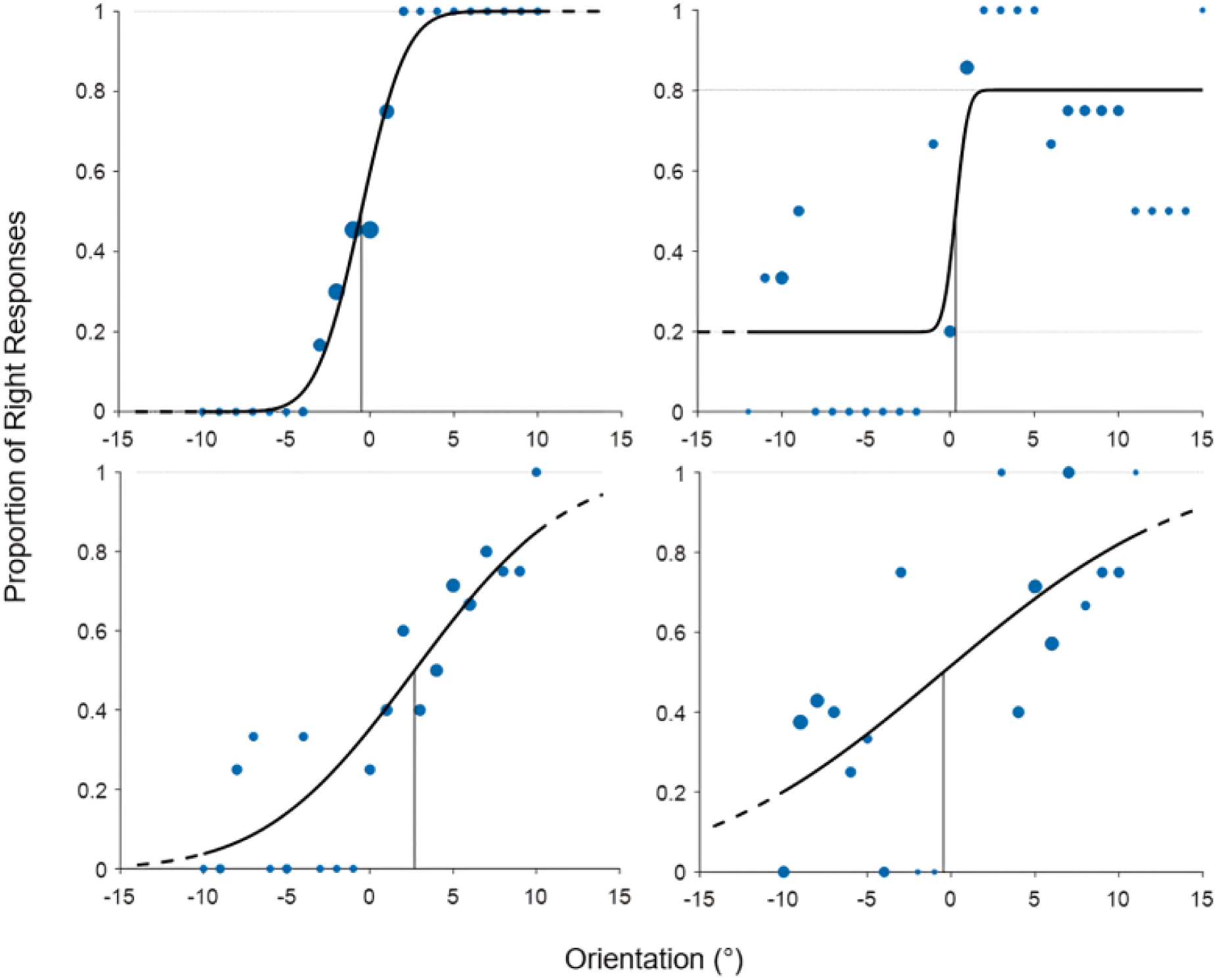
Examples of psychometric curves for good and poor task performance. The vertical lines represent the fitted thresholds, indicating the orientation perceived as vertical. Thin horizontal grey lines indicate the fitted lapse rates, which capture deviations from perfect performance at the extreme ends of the stimulus range. Blue dots show the mean proportion of right choices at each orientation, with dot area proportional to the square root of the number of trials at that angle. The transition from solid to dashed in the black curve corresponds to the range of the presented orientation values. The left panels show examples of acceptable fits: the top-left shows good performance with a steep slope and minimal lapse rate, and the bottom-left shows small deviations around negative to mid-range angles but no major misfits. Participants with these fits were included in the analysis. The right panels show poor fits: the top-right shows a high lapse rate (19.9%), and the bottom-right shows considerable deviations from the fitted curve across orientations. Participants with these fits were excluded.

To characterize overall patterns in participants’ decision-making performance, we examined the RTs of the decision response. We divided the data into twelve combinations defined by initial response (left/right), decision response (left/right), and stimulus onset delay (short, medium, long). For visualisation, we first computed each participant’s median RT within each of the twelve combinations and at each orientation, then averaged these medians across participants to yield group-level RT estimates. To formally test the RT differences related to initial response and stimulus onset delays while preserving trial-by-trial variability, we conducted an exploratory analysis using a linear mixed-effects model on trial-level RTs. The model included initial response, decision response, stimulus onset delay, and absolute orientation as fixed effects, with participants modelled as random intercepts.

To determine whether decision responses showed an alternation bias relative to initial responses across the three stimulus-onset delays, we compared the estimated threshold parameter (i.e., the orientation at which participants responded “right” 50% of the time) of the psychometric curve that was fitted for trials beginning with a left initial response and those with a right initial response. This method was adopted since the initial response was held constant within each block, making alternation probability an unsuitable measure for quantifying decision bias. For the fitted psychometric curves, a negative threshold indicates a rightward bias (less clockwise tilt required to choose “right”), while a positive threshold reflects a leftward bias (more clockwise tilt needed). For left-initial trials, an alternation bias would lead to a negative threshold whereas a repetition bias would lead to a positive threshold. The opposite pattern holds for right-initial trials: a negative threshold reflects repetition; a positive threshold reflects alternation. To obtain a single, hand-independent index of decision bias, we subtracted the right-initial threshold from the left-initial threshold values. Thus, negative decision bias indicates an alternation bias (a tendency to make the decision response opposite the initial response), while positive decision bias indicates repetition bias. We computed this choice bias value for each participant and delay, and tested it against zero using Wilcoxon signed-rank tests for each delay. Finally, we used the Friedman test to determine whether decision bias varied systematically across the three stimulus onset delays.

#### 2.5.2 EEG Analysis

##### 2.5.2.1 EEG preprocessing

Raw EEG data underwent preprocessing using the open-source toolbox FieldTrip (version 2024-07-04; Oostenveld et al., 2011) in MATLAB R2023b (The MathWorks, Inc.). To facilitate artifact rejection and ensure consistent independent component analysis (ICA), EEG data were initially segmented into long stimulus-locked epochs from –3 s to 8 s relative to the onset of the letter cue. As we were primarily interested in beta oscillations during the post-initial movement period and the decision-making phase, we subsequently time-locked the time series of each trial to the decision stimulus onset. A baseline interval was defined from –1.5 s to 0 s relative to the onset of the letter cue and was inferred from sample indices stored during the initial segmentation.

Epoched time series were first band-pass filtered between 0.1 and 70 Hz using a second-order bidirectional Butterworth filter, and a notch filter (49–51 Hz) was used to remove line noise. Artifact rejection was conducted in two stages. A first round of artifact rejection was performed based on visual inspection prior to ICA to eliminate prominent artifacts such as head movements that could impair ICA decomposition. Bad channels exhibiting excessive noise were identified and excluded during this step, which were then replaced with the average of their neighbouring channels. On average, 1.4 channels per participant were removed and replaced. ICA was then implemented to identify and remove ocular (e.g., blinks, saccades) and muscle artifacts while preserving beta-band features. On average, 21 components were rejected per participant (range: 9–27). A second round of visual inspection was then conducted to reject any residual artifacts. Prior to the second round of artifact rejection, the data were re-segmented for two stimulus-locked analyses: (1) epochs spanning –3.5 s to 3 s relative to the onset of the decision stimulus for event-related time-frequency analysis, and (2) epochs from –2.5 s to 1 s relative to the letter cue for baseline normalization. To ensure complete event capture and exclude abnormally slow initial responses to the letter cue, trials with initial response RTs exceeding 1.5 s were removed. Consistent with the behavioural analysis, trials were also excluded if the initial response was incorrect, or if no decision response occurred within the 3-s response window.

##### 2.5.2.2 Scalp**-**level time–frequency analysis

To inform source localization and virtual channel construction, we first performed scalp-level time–frequency analyses to identify beta-band desynchronization and rebound following initial responses. A sliding-window multi-taper Fourier transform was applied to two time-locked epochs: from –3.5 to 3 s relative to the grating onset, and from –2.5 to 1 s relative to the letter cue. Spectral power was estimated across frequencies from 5 to 40 Hz in 1 Hz steps, using 50 ms time steps and a frequency smoothing of 40% of the centre frequency (i.e. ±0.4 × f). Analyses were performed separately for each participant, initial response (left/right), decision response (left/right), and delay (short/medium/long). Power spectra time-locked to the decision stimulus onset were baseline-corrected using the –2.0 to 0 s window relative to the letter cue. Specifically, for each channel and frequency, power at each time point was expressed as a percentage change from the mean baseline power, computed by subtracting and then dividing by the baseline mean, and then multiplying by 100%. Grand-averaged, baseline-normalized beta-band activity (13–30 Hz) were used to define delay-specific time windows for subsequent source-level analyses: –0.75 to –0.25 s (desynchronization) and –0.25 to 0.25 s (rebound) for short delay; –1.5 to –1.0 s and –1.0 to –0.5 s for medium delay; and –3.0 to –2.75 s and –2.75 to –2.25 s for long delay, each relative to the decision stimulus onset. These windows corresponded to periods of strongest beta desynchronization and rebound induced by the initial response.

##### 2.5.2.3 Beamforming source localisation and seed extraction

The selected time windows were used to identify the source locations with the largest contrast between beta synchronization and desynchronization. For this, cross-spectral density matrices were calculated at 22 Hz with a spectral smoothing of ±8 Hz for each delay, initial response, and participant. A standard three-shell volume conduction model and a 1 cm resolution source grid in MNI space were used to construct forward models and compute lead field matrices. Frequency-domain beamforming was performed using the Dynamic Imaging of Coherent Sources (DICS) method (Gross et al., 2001). A common spatial filter was computed from the combined time windows and then applied separately to each window to estimate source-level power. Beta band power was computed as the normalised difference between the rebound and desynchronization windows, by dividing their power difference by the average power across the two windows. The resulting source-level beta activity maps were averaged across delays and initial responses. To identify motor-related regions, left hemisphere activity was examined using right initial-response trials, and right hemisphere activity using left initial-response trials. The motor-related anatomical regions were identified using the SPM Anatomy Toolbox (Eickhoff et al., 2005). Group-level peak coordinates were identified in each hemisphere, and individual peaks were selected within a 3 cm radius of these group-level maxima. These individual peak coordinates served as seed points for virtual channel extraction, referred to as the left motor cortex (LMC) and right motor cortex (RMC). Source-level time series were extracted using the linearly constrained minimum variance (LCMV) beamforming method based on covariance matrices that were computed for the entire –3.5 to +3 s time window relative to decision stimulus onset and frequencies between 5 and 40 Hz. The resulting spatial filters were then applied to the scalp-level data to extract single-trial virtual channel time series for each seed. This procedure was repeated for the epoch from –2.5 to 1 s time-locked to the letter cue.

##### 2.5.2.4 Source-level time-frequency analyses and statistical testing

Source-level time–frequency representations (TFRs) of the virtual channel signals were computed using a single-taper sliding-window Fourier transform with a Hanning taper (5–40 Hz range in 1 Hz steps, 50 ms timesteps, 5 cycles). TFRs were computed separately for LMC and RMC, initial response (left/right), decision response (left/right), and delay (short, medium, long). Power spectra time-locked to the decision stimulus were baseline-corrected using the –2 to 0 s interval from the letter cue-locked data, by calculating the percentage change from the mean baseline power. The resulting grand-averaged, normalized power spectra were visualized across all conditions and seed locations.

To investigate whether lateralized beta power at the onset of the decision period varied across stimulus onset delays, we extracted beta power (13–30 Hz) within the 0–0.25 s window during which the decision stimulus was displayed. For each trial, beta power was computed separately for the motor cortex contralateral and ipsilateral to the initial motor response. We then calculated the difference between contralateral and ipsilateral beta power (hereafter referred to as contra–ipsi beta), providing a measure of lateralized sensorimotor beta power. The values were averaged across the defined time and frequency window, and a one-way repeated-measures ANOVA was conducted to test for a significant effect of stimulus onset delay on contra–ipsi beta.

#### 2.5.3 Brain–Behaviour Correlations

To assess whether stronger lateralized sensorimotor beta power during decision stimulus presentation indeed predicts alternation bias, we computed one-tailed Spearman’s rank correlations both across and within subjects. We focused these analyses on the most difficult trials with orientation values constrained between -5° and +5°, as choice history biases are known to be strongest when perceptual evidence is weak. Prior work (Fründ et al., 2014) showed that up to 32% of decision variance on difficult trials (55–75% accuracy) is attributable to prior choices, whereas on easier trials (>75% accuracy), behaviour is almost entirely stimulus-driven. Across subjects, we correlated each participant’s mean contra–ipsi beta power (averaged across the three onset delay conditions) with their mean choice bias, testing the a priori hypothesis that individuals with higher beta lateralization would exhibit stronger alternation bias (i.e., more negative bias values). To eliminate between-subject variability in beta power and choice bias, we also performed within-subject correlations. Specifically, we computed residual values by subtracting each participant’s mean contra–ipsi beta power and choice bias from their respective values in each delay condition. We then applied one-tailed Spearman’s rank correlations to these residuals to test whether delay-specific fluctuations in beta power predicted stronger alternation bias. For exploratory purposes, we further examined whether lateralized sensorimotor beta power is associated with decision speed by conducting the same across- and within-subject analyses, using two-tailed Spearman’s rank correlations between beta power and RTs. To assess whether this relationship depends on the congruency between the initial and decision responses, given the inhibitory role of beta activity in suppressing repeated actions, we further split the data into repeating and alternating trials and computed two-tailed Spearman correlations separately for each subset.

## 3. Results

### 3.1 Behavioural Results

We firstly characterised the overall behavioural pattern in participants’ decision-making performance using the median RTs across combinations of initial response (left/right), decision response (left/right) and stimulus onset delay (short/medium/long). Median RTs were computed per participant for each combination and orientation and then averaged across participants. As shown in **Figure 5**, RTs exhibited a consistent trend: responses were slower for grating stimuli near the vertical axis, in line with increased task difficulty at more vertical orientations. Notably, across all three stimulus onset delays, participants appeared to respond consistently faster following right initial responses compared to left initial responses. This observation was supported by an exploratory trial-level linear mixed-effects analysis, which revealed that trials with right initial responses (median = 0.549 s, IQR = [0.441, 0.690] s) were significantly faster than those with left initial responses (median = 0.573 s, IQR = [0.463, 0.727] s), β = –0.0506 s, SE = 0.0043 s, *t* = –11.66, *p* < 0.001. Additionally, trials with right decision responses (median = 0.552 s, IQR = [0.450, 0.695] s) were significantly faster than those with left decision responses (median = 0.569 s, IQR = [0.452, 0.717] s), β = –0.0178 s, SE = 0.0043 s, *t* = –4.11, *p* < 0.001. RTs were also influenced by stimulus onset delays: the short delay (median = 0.583 s, IQR = [0.469, 0.728] s) resulted in significantly slower responses compared to the medium (median = 0.544 s, IQR = [0.442, 0.691] s) and long delay (median = 0.552 s, IQR = [0.446, 0.699] s). Longer delays significantly accelerated decision speed (medium: β = –0.0395 s, SE = 0.0053 s, *t* = –7.41, *p* < 0.001; long: β = –0.0299 s, SE = 0.0053 s, *t* = –5.62, *p* < 0.001), as illustrated in **Figure 6A**.

**Figure 5.**
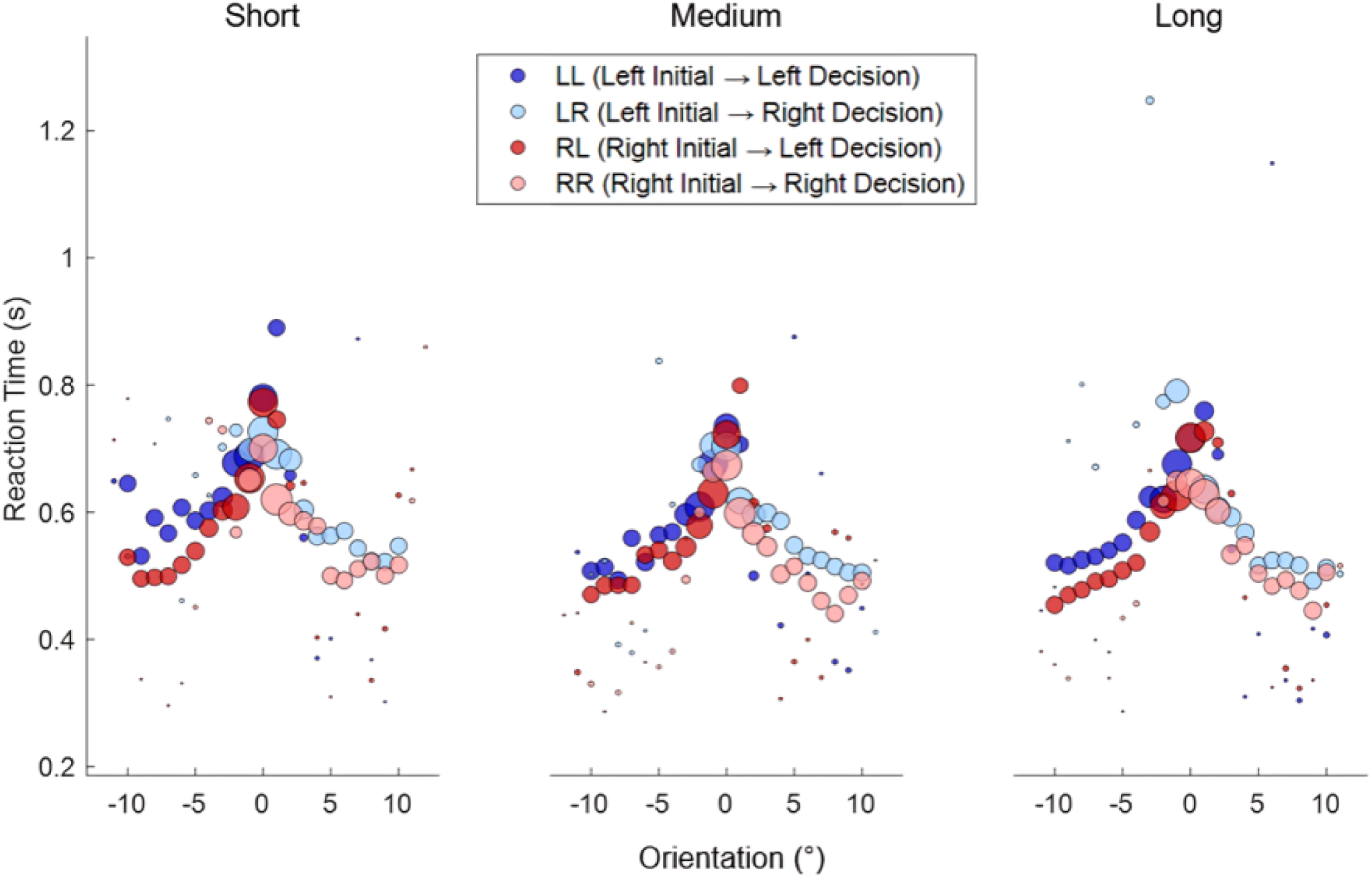
Group-level median RTs as a function of stimulus orientation. Each subplot combines initial (left/right) and decision (left/right) response types and displays data per stimulus onset delay (short, medium, long). Positive orientation values represent clockwise stimuli, while negative values indicate anticlockwise stimuli. Task difficulty increases as orientation approaches 0°, corresponding to vertical. The area of each data point represents the proportion of trials associated with each orientation and combination.

**Figure 6.**
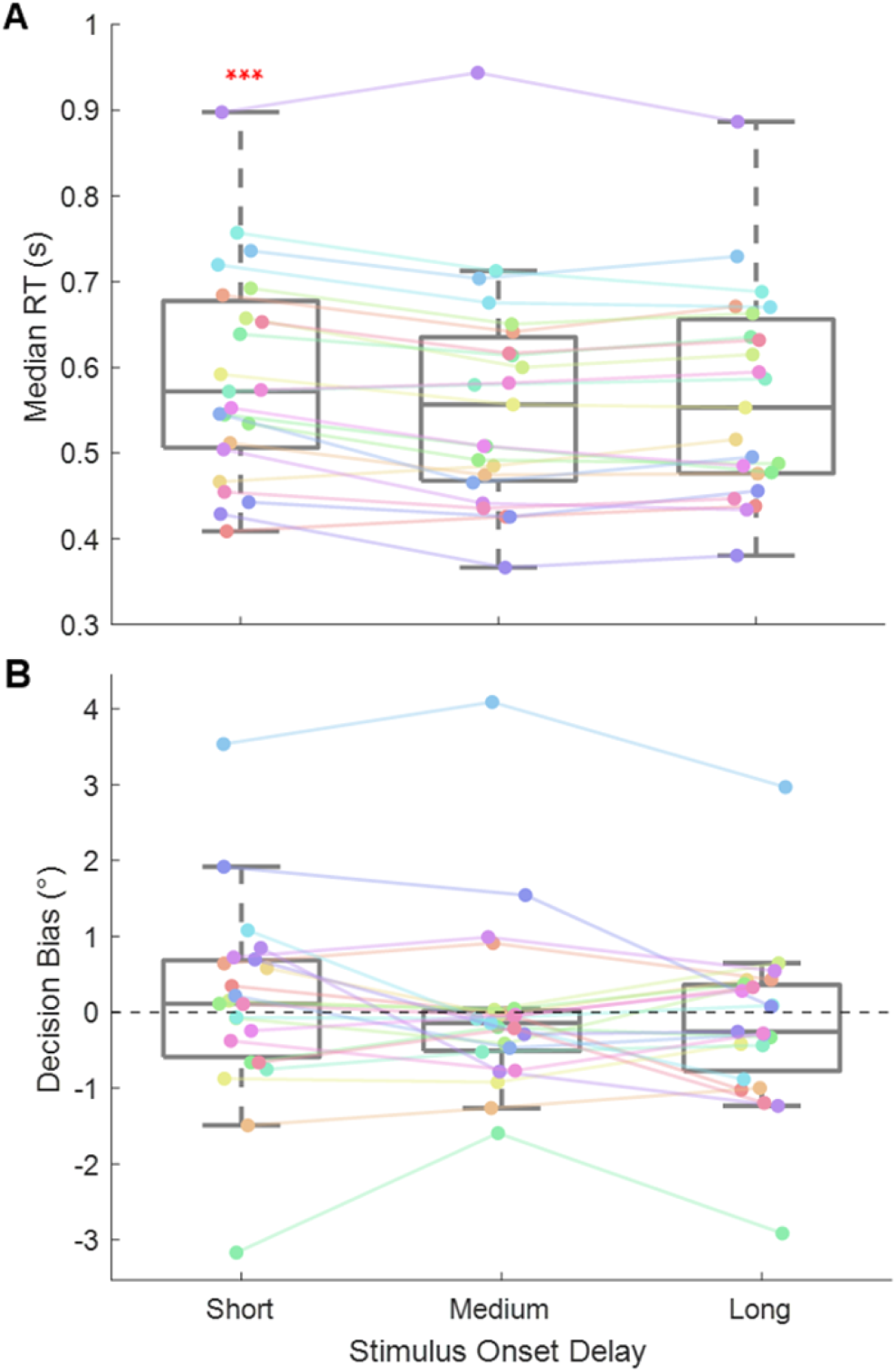
**A.** Individual participants’ median reaction times across stimulus onset delays. In each boxplot, the central line represents the median, while the upper and lower edges indicate the 75th and 25th percentiles, respectively. Whiskers indicate the most extreme values within 1.5 times the interquartile range. Individual data points are connected by coloured lines to illustrate within-subject trajectories across the three delays. Three red asterisks indicate *p* < .001, suggesting a significant slower RTs observed in the short delay compared to the medium and long delay. **B.** Decision bias computed as the threshold difference between left- and right-initial response trials, plotted across the three stimulus onset delays. Negative values indicate an alternation bias (i.e., a tendency to make a decision opposite to the initial response), while positive values reflect a repetition bias. Boxplots follow the same format as in (A), with coloured lines connecting each participant’s data across delays.

To further examine whether the initial response biased subsequent decisions towards alternation, we fitted individual psychometric functions for trials beginning with a left versus a right initial response at each stimulus onset delay. We quantified decision bias as the difference in decision thresholds (i.e., the orientation at which participants responded “right” 50% of the time) between left-initial and right-initial trials. Negative decision bias indicates an alternation bias (i.e., a tendency to make a decision opposite to the initial response), while positive bias reflects a repetition bias. As shown in **Figure 6B**, a positive mean decision bias was observed in the short delay (0.11°, 95% CI = [–0.40°, 0.62°]), indicating that the threshold for left-initial trials (Mean = –0.19°, SD = 1.02°) was higher than that for right-initial trials (Mean = –0.25°, SD = 0.85°). This result suggested a tendency toward repetition through a higher likelihood of making a decision consistent with the initial response. However, the effect was not statistically significant, as revealed by a Wilcoxon signed-rank test (Mdn = 0.11°, *W* = 156, *z* = 0.55, *p* = 0.584). In contrast to the short delay, negative decision biases were observed in both the medium (Mean = –0.01°, 95% CI = [–0.47°, 0.45°]) and long delay (Mean = –0.16°, 95% CI = [–0.60°, 0.27°]). In the medium delay, thresholds were slightly lower following left-initial responses (Mean = –0.30°, SD = 1.01°) than right-initial responses (Mean = –0.29°, SD = 0.84°), suggesting a weak alternation bias. A similar pattern emerged in the long delay, where thresholds following left-initial responses (M = –0.16°, SD = 1.06°) were lower than those following right-initial responses (Mean = 0.00°, SD = 0.72°). However, neither effect reached statistical significance (medium: Mdn = –0.14°, *W* = 87, *z* = –1.55, *p* = 0.121; long: Mdn = –0.26°, *W* = 112, *z* = –0.79, *p* = 0.429). These findings suggest that initial motor responses do not exert a systematic bias on subsequent perceptual decisions at a group level. Moreover, decision bias did not significantly differ between the three stimulus onset delays, as indicated by a Friedman test, χ²(2, N = 23) = 1.04, *p* = .594, Kendall’s W = .02. The absence of a significant difference may be attributed to substantial individual variability (consistent across the three delays), with some participants exhibiting alternation biases and others showing repetition biases.

### 3.2 EEG Results

We identified the source locations with the largest contrast between beta synchronization and desynchronization induced by the initial response in order to extract source-level time series. Beamformer source images were computed for each participant, type of initial response and stimulus onset delay, and subsequently grand-averaged separately for left and right initial responses. Results reveal prominent beta activity in the hemisphere contralateral to the initial response hand (**Figure 7**). Group-level peak coordinates were identified in each hemisphere. In the left hemisphere (based on right initial-response trials), peak beta activity was localized at MNI coordinates [–40, 0, 60] mm, corresponding to the left premotor and middle frontal gyrus. In the right hemisphere (based on left initial-response trials), the peak was observed at [40, –20, 70] mm, localizing to the right primary motor cortex. We refer to these locations as LMC and RMC in the remainder of the manuscript.

**Figure 7.**
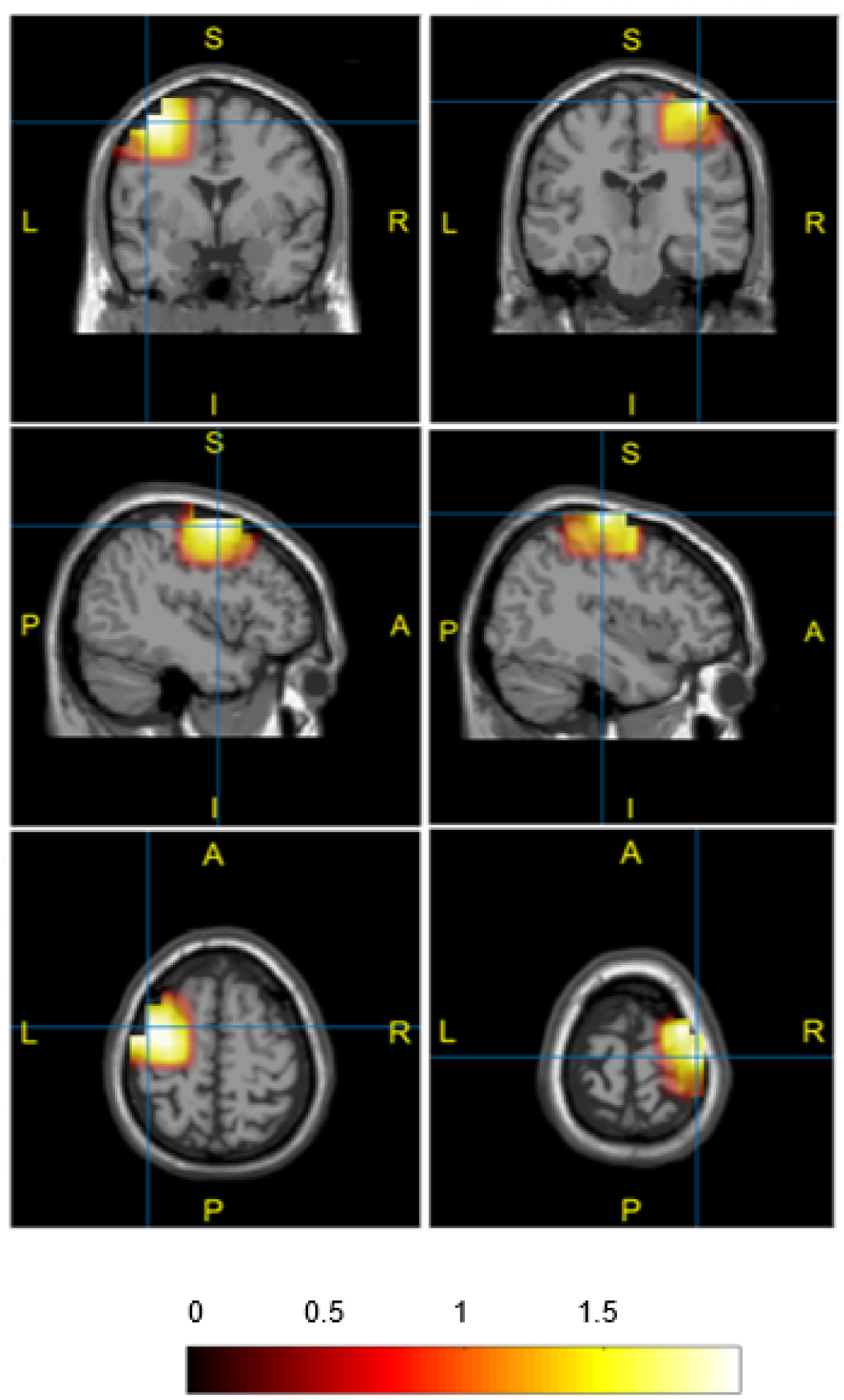
Grand-averaged source maps of the contrast between beta-band rebound and desynchronization induced by the initial response. In the left panels (right initial-response trials, assessing left-hemisphere activity), the peak beta activity was localized at MNI coordinates [–40, 0, 60] mm, consistent with the left premotor and middle frontal gyrus. In the right panels (left initial-response trials, assessing right-hemisphere activity), the peak was observed at [40, –20, 70] mm, within the right primary motor cortex. The color bar reflects the magnitude of the normalised beta power difference, with positive values indicating stronger beta rebound related to desynchronization. Individualized peak coordinates were identified within a 3 cm radius of the group-level peaks and served as the LMC and RMC seed locations for subsequent virtual channel extraction.

Grand-averaged, baseline-corrected source-level TFRs for LMC and RMC across all combinations of initial response (left/right) and decision response (left/right), for the short, medium, and long stimulus onset delay conditions are shown in **Figures 8–10**. Across all onset delay conditions, beta power was still decreased for approximately 250 ms following the termination of the initial response, followed by a prominent beta rebound in the hemisphere contralateral to the responding hand, lasting approximately 500 ms. Following the onset of the decision stimulus, a second beta decrease emerged rapidly, lasting until approximately 250 ms after the initiation of the decision response. This was again followed by a robust beta rebound lasting for approximately 1 s. The second beta rebound was stronger contralateral to the decision response hand in trials with left initial responses, as expected. However, in trials beginning with right initial responses, the rebound remained consistently stronger over the LMC, regardless of the expected lateralization.

**Figure 8.**
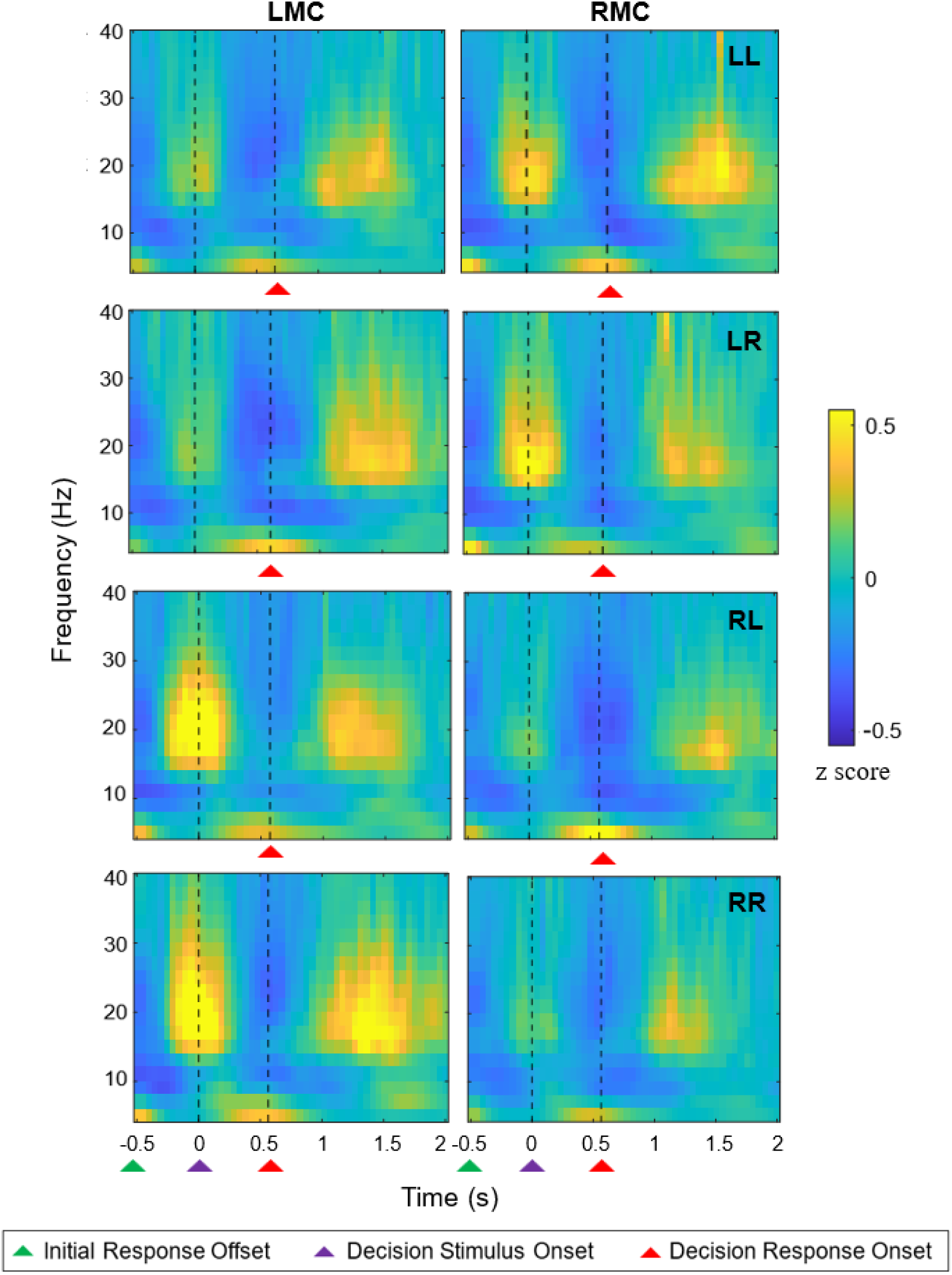
Grand-averaged time-frequency spectra for the LMC and RMC across all combinations of initial response (left/right) and decision response (left/right) for the short onset delay. Power spectra are time-locked to the onset of the decision stimulus and baseline-corrected using the –2 to 0 s interval relative to the letter cue. Each horizontal pair of panels represents one initial–decision response combination (e.g., “LL” = left initial response and left decision response), with LMC activity shown on the left and RMC activity on the right. The green triangles indicate the termination of the initial response. The purple triangles indicate the onset of the decision stimulus presentation. The red triangles indicate the median RT of the decision response.

**Figure 9.**
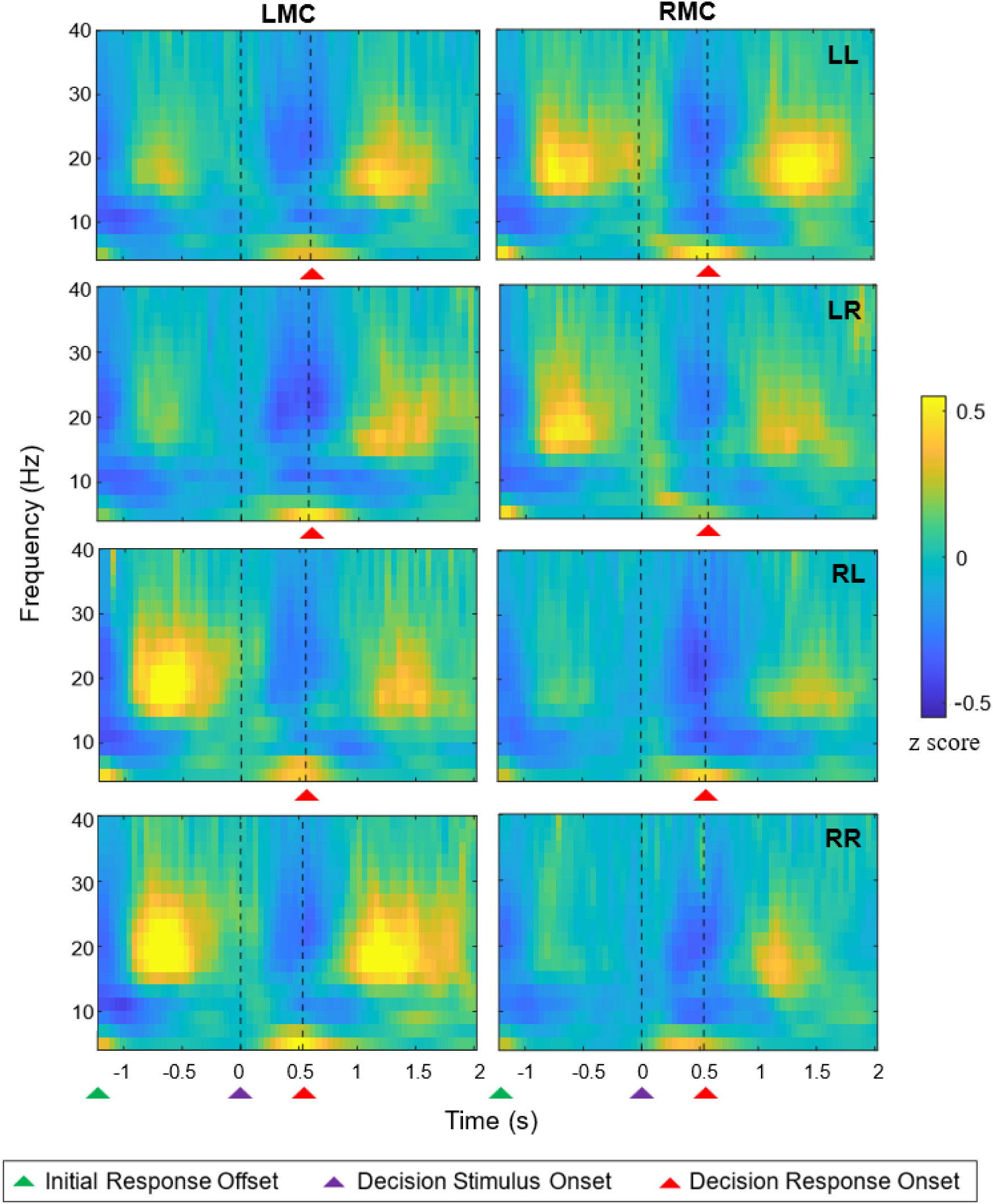
Grand-averaged time-frequency spectra for the LMC and RMC across all combinations of initial response (left/right) and decision response (left/right) for the medium onset delay. Additional details are provided in Figure 8.

**Figure 10.**
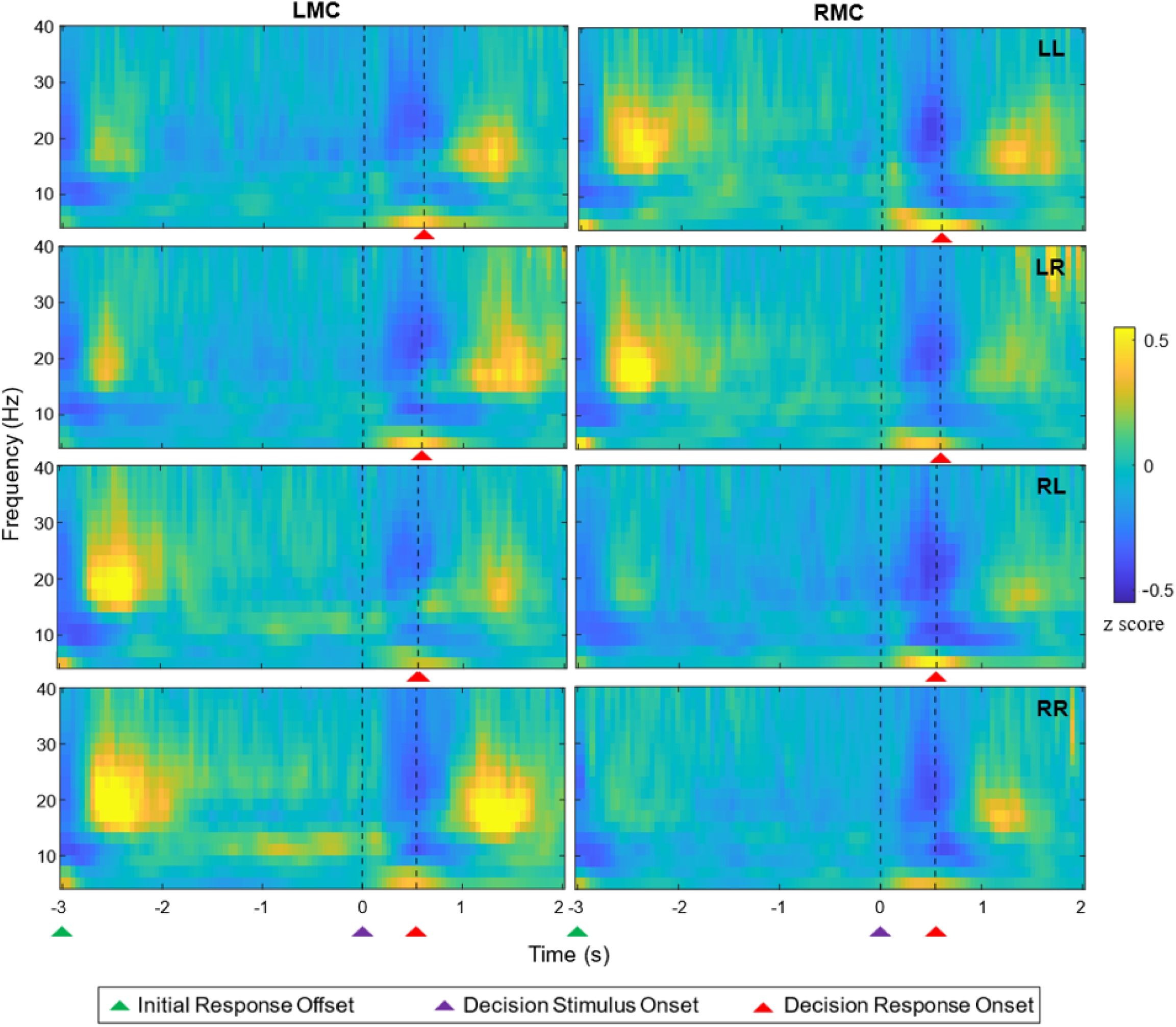
Grand-averaged time-frequency spectra for the LMC and RMC across all combinations of initial response (left/right) and decision response (left/right) for the long onset delay. Additional details are provided in Figure 8.

To better visualize and compare the temporal dynamics of lateralized beta power around decision onset across the three delay conditions, we computed contra–ipsi beta power time series relative to the initial response hand for each delay, as shown in **Figure 11**. Contra–ipsi beta power was clearly elevated at the time of decision stimulus onset in the short delay condition, partially persisted in the medium delay, and returned more fully to baseline in the long delay condition. This pattern reflected the significant main effect of delay, as revealed by a repeated-measures ANOVA on contra–ipsi beta power averaged across 13–30 Hz and the 0–250 ms window of decision stimulus presentation (**Figure 12**). Mean contra–ipsi beta power was highest in the short delay condition (Mean = 0.243, SD = 0.231), lower in the medium delay (Mean = 0.149, SD = 0.087), and lowest in the long delay condition (Mean = 0.087, SD = 0.106), *F*(2, 44) = 6.72, *p* = .003, η² = .23 (Greenhouse–Geisser corrected *p* = .011). Bonferroni-adjusted post hoc comparisons revealed significantly lower contra–ipsi beta power in the long delay condition compared to both the medium delay (mean difference = –0.062, SE = 0.021, *p* = .021) and short delay (mean difference = –0.156, SE = 0.053, *p* = .024). The difference between the short and medium delay conditions was not statistically significant (mean difference = –0.094, SE = 0.047, *p* = .170).

**Figure 11.**
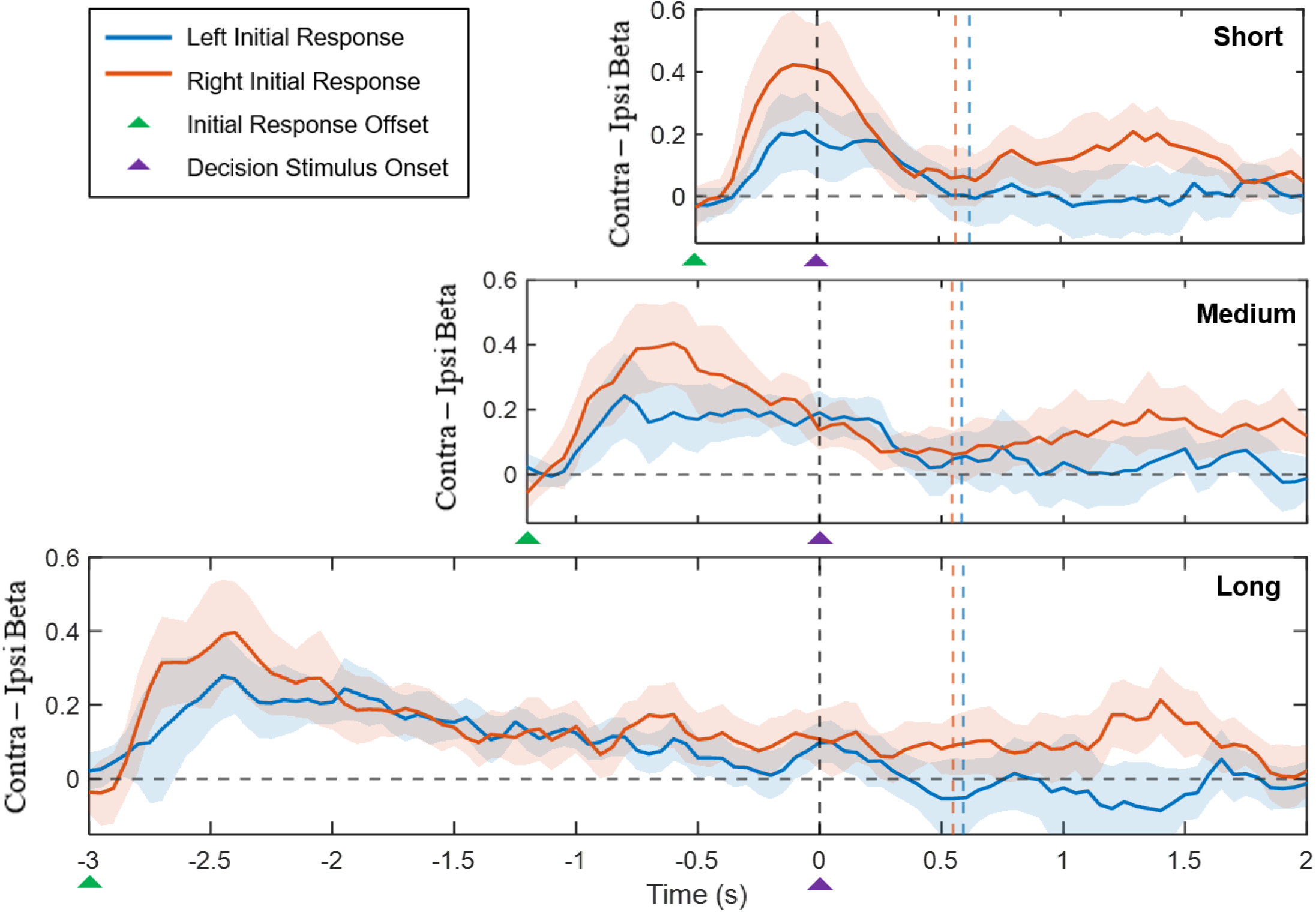
Lateralized beta power time series relative to the initial response hand across short, medium, and long delay conditions (top to bottom panels), time-locked to the onset of the decision stimulus and baseline-corrected using the –2 to 0 s interval relative to the letter cue. The blue solid line shows beta power for left initial-response trials, computed as RMC minus LMC; the blue dashed vertical line indicates the median RT for the decision response in these trials. The red solid line shows beta power for right initial-response trials, calculated as LMC minus RMC; the red dashed vertical line marks the median RT for the decision response in these trials. The shaded areas indicate the 95% confidence interval of the mean lateralized beta time course.

**Figure 12.**
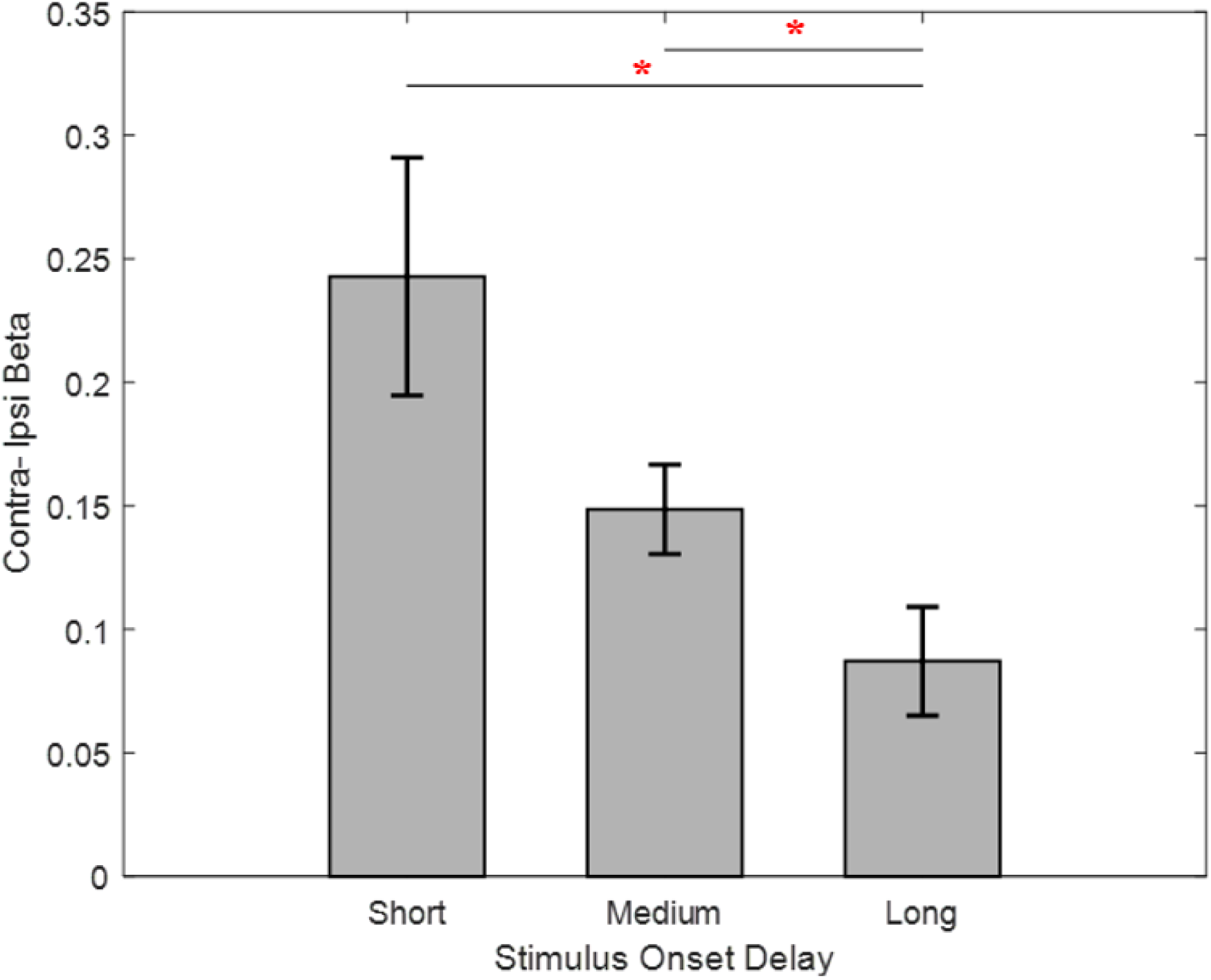
Lateralized beta power averaged across the 13–30 Hz beta frequency range and the 0–250 ms window of decision stimulus presentation, shown for the short, medium, and long delay conditions. Bar heights represent the mean value across participants, and whiskers indicate the standard error of the mean (SEM). Contra–ipsi beta power decreased progressively with increasing delay duration. Significant differences were observed between the short and long delay conditions (*p* = .024), as well as between the medium and long conditions (*p* = .021), as determined by repeated-measures ANOVA with Bonferroni-adjusted post hoc tests. The red asterisk indicates *p* < .05.

### 3.3 Brain-Behaviour Correlations

To test whether stronger beta lateralization at the onset of decision-making biases subsequent decision responses toward alternation, we first examined the between-subject correlation between mean contra–ipsi beta power and mean decision bias, averaged across the three onset delays. A one-tailed Spearman’s rank correlation tested the a priori hypothesis that higher beta lateralization would predict stronger alternation bias (i.e., more negative bias values). In line with our hypothesis, we found a significant negative correlation (ρ = –0.366, one-tailed *p* = .043; **Figure 13A**), indicating that individuals with stronger beta lateralization at decision onset were more likely to alternate their decision responses relative to their initial responses. At the within-subject level, we tested whether delay-specific variations in contra–ipsi beta power predicted corresponding changes in decision bias. For each participant, residual values (delay-specific values minus the participant’s mean across delays) were computed, yielding three data points per individual. A one-tailed Spearman’s rank correlation again revealed a negative but this time non-significant association (ρ = –0.155, one-tailed *p* = .102), indicating that while stronger beta lateralization tended to predict increased alternation bias within individuals, the effect was not strong enough to significantly explain variations in alternation bias across delays.

**Figure 13.**
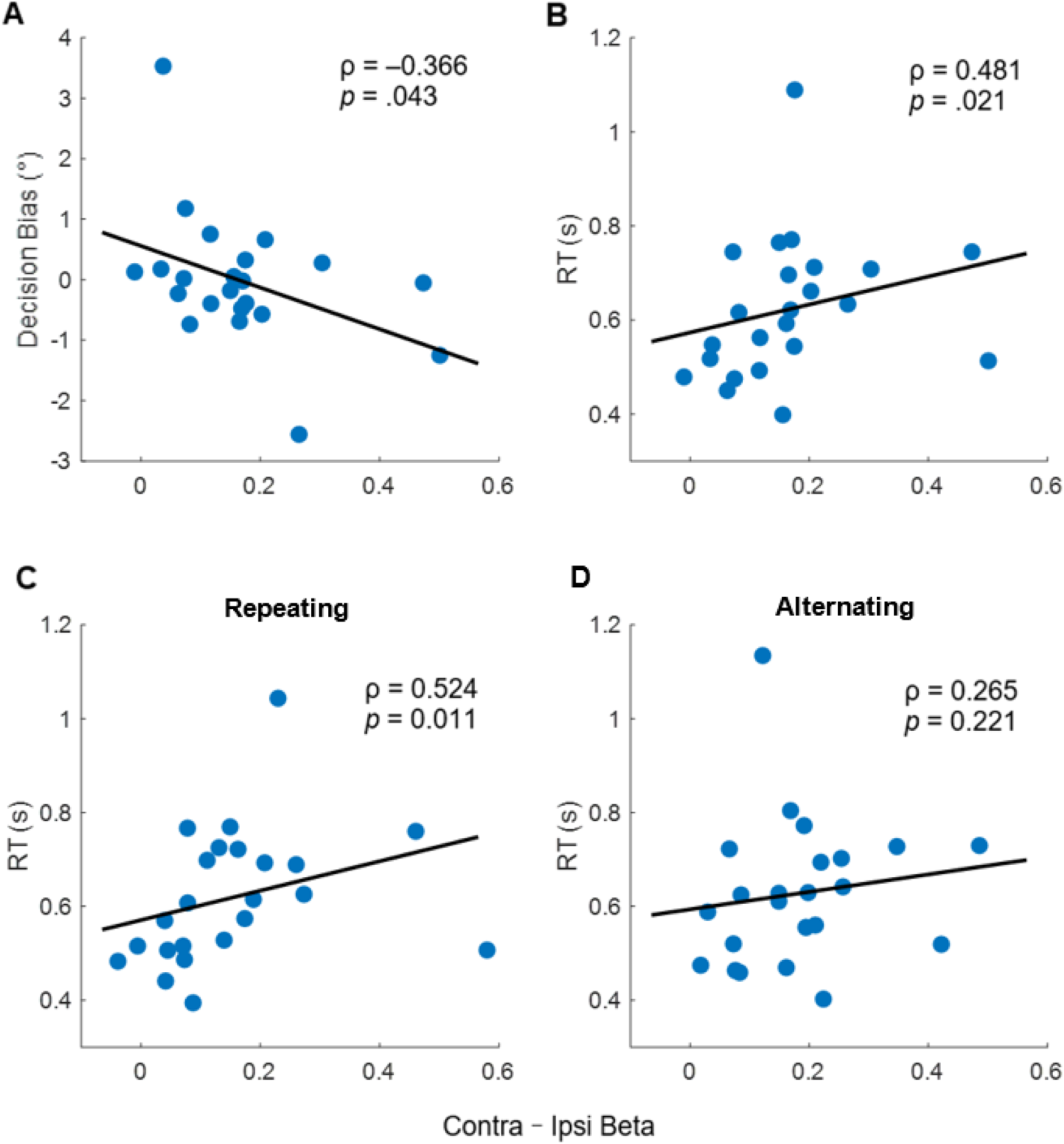
Between-subject Spearman correlations between contra–ipsi beta power and behavioral measures averaged across the three onset delays. **(A)** Negative correlation between contra–ipsi beta power and decision bias, indicating that greater beta lateralization was associated with stronger alternation bias. **(B)** Positive correlation between contra–ipsi beta power and RTs, suggesting that higher beta power predicted slower decision speed. Correlations between beta power and RTs shown separately for repeating trials **(C**), where the initial and decision responses were made on the same side, and alternating trials (**D**), where the two responses occurred on opposite sides. Each dot represents one participant’s mean value averaged across all three onset delays.

Finally, we conducted a set of two-tailed Spearman’s rank correlations between contra–ipsi beta power and RTs as an exploratory analysis to test whether sensorimotor beta oscillations predicted decision speed. At the between-subject level, higher mean contra–ipsi beta power was significantly associated with slower RTs (ρ = 0.481, *p* = .021; **Figure 13B**), indicating that individuals with stronger beta lateralization around decision onset tended to respond more slowly. This positive relationship between beta power and RTs was significant only in repeating trials, where the initial response was made on the same side as the decision response (ρ = 0.524, *p* = .011; **Figure 13C**), but not in alternating trials, where the two responses occurred on opposite sides (ρ = 0.265, *p* = .221; **Figure 13D**). No significant within-subject correlation was observed between delay-specific variations in beta power and RTs (ρ = –0.030, *p* = .724), indicating no significant correlation between beta lateralization and decision speed within individuals.

## 4. Discussion

The aim of the present study was to investigate whether prior motor states can bias subsequent perceptual decisions through the lens of sensorimotor beta oscillations. We designed a perceptual decision-making task in which participants judged whether a briefly presented orientation grating was tilted clockwise or anticlockwise by responding with a left- or right-hand button press. Each decision was preceded by an initial, choice-unrelated motor action, cued by letters and requiring a left- or right-hand button press, with three variable delays separating this initial motor response and decision stimulus presentation. This design allowed us to probe whether the temporal dynamics of beta oscillations, evoked by the initial motor response and lateralized in strength between left and right sensorimotor cortices, could influence subsequent perceptual decisions and reaction times. Our study addresses possible neural mechanisms of embodied decision-making by investigating whether sensorimotor beta activity plays an active role in shaping choice formation.

Behaviourally, we found no systematic decision bias as a function of the initial motor response, indicated by the lack of a significant difference in threshold for the psychometric functions fitted separately to trials following left versus right initial responses. This suggests that prior motor activity did not influence subsequent perceptual decisions at the group-level. This finding is consistent with those of Akaishi et al. (2014) and Braun et al. (2018), both of whom used variable stimulus-response mappings and unpredictable delays, and found that sequential dependencies in behaviour were attributed primarily to previous choices, rather than to motor history. By contrast, Pape et al. (2017) and Zhang & Alais (2020) reported clear motor history effects. Zhang & Alais (2020) observed a motor alternation bias under variable stimulus-response mappings, indicating that motor history can influence decisions independently of perceptual choice history. Pape et al. (2017), using a similar paradigm to ours with a choice-unrelated motor action and variable response mappings, found an increased tendency to alternate choices. These discrepancies in results may reflect differences in task structure, as also observed by Fründ et al. (2014). For example, the inclusion of variable delays, as in our study and those by Akaishi et al. (2014) and Braun et al. (2018) may interrupt automatic response tendencies and attenuate motor history effects. Moreover, the presence or absence of feedback can impact decision strategies, such that feedback may prompt participants to shift their decision criterion following errors (Fründ et al., 2014; Pape et al., 2017). Task difficulty is another factor: choice-history biases are often more pronounced in difficult trials when stimulus evidence is weak, and decisions rely more on internal priors (Fründ et al., 2014). In addition, decision uncertainty has been shown to modulate history biases, either dampening or amplifying them depending on context (Braun et al., 2018; Koizumi et al., 2015; Urai et al., 2017). Our use of a choice-unrelated initial action, multiple delays, and group-level comparisons based on psychometric function threshold shifts rather than trial-level modelling, may have reduced the sensitivity to detect subtle motor history effects at the behavioural level.

Another possibility is that susceptibility to decision bias might reflect an individual trait, with some participants showing alternation tendencies and others showing repetition, potentially cancelling out any group-level effect (Fründ et al., 2014; Urai et al., 2019; Urai & Donner, 2022). Such inter-individual differences seem to be reflected in our neurophysiological data: participants with stronger beta power lateralization around decision onset exhibited a greater tendency to alternate their initial responses. This effect is likely driven by the inhibitory role of beta oscillations (Jurkiewicz et al., 2006; Pfurtscheller et al., 1996; Van Wijk et al., 2009, 2012), which may suppress the repetition of recently executed actions. These findings suggest that beta lateralization may serve as a sensitive marker of motor cortex involvement in decision-making, revealing individual differences in decision strategies that are not necessarily captured by group-mean behaviour under the current experimental conditions. Pape & Siegel (2016) also reported between-subject associations between beta lateralization and response alternation, albeit with stronger effects than those observed in our study. This might be due to the different approaches used to quantify decision bias: whereas we employed psychometric curve threshold shifts which provide mechanistic insight into overall decision bias while controlling for stimulus intensity, Pape & Siegel (2016) used trial-by-trial alternation rates, which primarily capture sequential switching tendencies. Additionally, the extended delay we introduced between the initial motor response and the decision stimulus presentation may have attenuated the influence of motor history on subsequent choices. Given that response alternation tendencies are most pronounced in fast response trials that immediately follow previous motor actions (Leite & Ratcliff, 2011; Urai et al., 2019; White & Poldrack, 2014), this effect was likely diminished in our design due to the inserted temporal delay.

In contrast to the observed between-subject association, we failed to detect significant within-subject correlations between beta lateralization and decision bias, suggesting delay-specific fluctuations in beta power did not reliably predict corresponding changes in decision bias within individuals. This contrasts with findings from Pape & Siegel (2016), who reported significant single-trial correlations. However, such trial-level analyses were not feasible in our study due to the use of psychometric functions, which fit to a set of trials per condition rather than providing single-trial measures. Urai and Donner (2022) also failed to detect a mediating effect of beta-band motor history signals on choice alternation. However, by applying single-trial modeling, they were able to show that motor history signals shift the starting point of the decision process toward alternation, revealing a subtle trial-by-trial modulation which was not evident at the condition level. Although our group-level analysis revealed the strongest beta lateralization around decision onset for the shortest delay, the precise timing and magnitude of beta activity may have varied substantially across delays within individuals, consistent with prior evidence that motor beta activity is highly dynamic and transient on a trial-by-trial basis (Little et al., 2019). Furthermore, participants may have adopted different decision strategies depending on the duration of the delay preceding the decision stimulus such as varying levels of anticipation and attention, potentially introducing confounding factors at the trial level.

Our exploratory analyses also examined how prior motor states influence subsequent decision speed. Behaviourally, we found that executing right-hand responses, regardless of whether they occurred during the initial action or decision phase, consistently led to faster reaction times. Given that the majority of our participants were right-handed, this finding aligns with previous studies reporting a response speed advantage for the dominant hand (Dexheimer et al., 2022; Kerr et al., 1963). This dominant-hand advantage is possibly due to the larger motor representation for the dominant limb (Triggs et al., 1994; 1999) and lower thresholds for motor cortical excitability in the left hemisphere (Macdonell et al., 1991; Triggs et al., 1994). Beyond faster motor preparation, this advantage might also reflect the dynamics of the decision process itself that the brain can favor right-hand responses even before sensory evidence is accumulated. At the neurophysiological level, participants with stronger beta lateralization tended to exhibit slower decision responses, an effect only observed in repeating trials where the initial and decision responses were made with the same hand. This finding can again be interpreted with the inhibitory role of beta oscillations (Jurkiewicz et al., 2006; Pfurtscheller et al., 1996; van Wijk et al., 2009, 2012), and is consistent with previous research showing that increased post-movement beta rebound predicts longer preparation times for subsequent actions (Muralidharan & Aron, 2021), likely mediated by GABAergic inhibitory mechanisms within the motor system (Zhang et al., 2024). Consistent with our findings on decision bias, we also failed to detect within-subject correlations between variations in beta lateralization and decision speed across delays. This may again suggest that beta activity represents an individual characteristic (Urai et al., 2019; Urai & Donner; 2022), while its temporal dynamics may vary flexibly across the delays used in our design.

Another interesting observation is that the beta rebound around the decision phase appeared more bilateral compared to the rebound following the initial motor response (see Figure 8-10). Bilateral beta rebound has been previously reported in both voluntary movement (Heinrichs-Graham et al., 2017; Jurkiewicz et al., 2006) and decision-making tasks (Donner et al., 2009; Pape & Siegel, 2016). This pattern could arise either from a carryover effect of the initial motor response especially when the time delay is short, or from the broader integration of associative regions during decision-making, which engages bilateral cortical networks more than simple motor execution. Furthermore, on trials that began with right-hand responses, the beta rebound remained consistently stronger over the left motor cortex, likely because the rebound threshold is lower in the contralateral (left) motor cortex for right-handed subjects, which allows left-hemispheric beta to dominate even during the subsequent decision phase.

Several limitations of the present study should be noted. To isolate motor-induced decision bias, we fixed the initial motor response cue within each block, ensuring voluntary movement without decision demands. While this design ensures a clean manipulation of motor cortex activity, it oversimplifies the interactive nature of decision-making, which involves continuous communication between motor and associative regions (e.g., prefrontal and parietal cortex). Additionally, fitting psychometric curves allowed us to compute overall decision bias, but it lacks sensitivity at the single-trial level, potentially overlooking finer sequential dynamics. The inclusion of three different decision stimulus delays appeared to shift individual beta dynamics in idiosyncratic ways. Future work should therefore explore how more fine-grained features of beta oscillations such as beta burst timing modulate decision behaviour in individual trials. Finally, our findings also invite exploration of oscillatory markers in the motor cortex beyond beta activity. Prior studies have highlighted that sensorimotor alpha-band oscillations can build up prior to a response and may encode error decisions (Urai & Donner, 2022) or predict decision biases (Zhang et al., 2019). Whether this alpha modulation reflects top-down regulation of decision formation or simply motor preparation remains future investigation.

In summary, we investigated how prior motor states influence subsequent decision bias by embedding an initial left- or right-hand movement with variable delays before a perceptual decision-making task. Although the initial movement did not lead to a significant group-level decision bias or consistent within-subject correlations between decision bias and lateralized beta power across delays, individuals with stronger beta lateralization around decision onset showed greater alternation tendencies and slower decision speeds. Following the perspective of embodied-decision making, our results partially support the influence of motor cortex activity on the choice process, while also suggesting that beta activity may serve as a trait-like index of individual susceptibility to decision bias.

